# Synapse propensity of human memory CD8 T cells confers competitive advantage over naïve counterparts

**DOI:** 10.1101/507392

**Authors:** Viveka Mayya, Edward Judokusumo, Enas Abu-Shah, Willie Neiswanger, Lance C Kam, Michael L Dustin

## Abstract

Memory T cells are endowed with multiple functional features that enable them to be more protective than naïve T cells against infectious threats. It is not known if memory cells have a higher synapse propensity, i.e. increased probability to form immature immunological synapses that then provide an entry into different modes of durable interaction with antigen presenting cells. Here we show that only human memory CD8 T cells have remarkably high synapse propensity compared to naïve counterparts. Such a dichotomy between naïve and memory cells is not observed within the human CD4 or murine CD8 T cell population. Increased surface expression of LFA1 contributes to the higher synapse propensity in human memory CD8 T cells. Finally, we show that higher synapse propensity in human memory CD8 T cells allows them to compete out naïve CD8 T cells from getting recruited to the response. This observation has implications for original antigenic sin and aging of the immune system in humans.

## Introduction

Memory T cells exhibit functional avidity maturation that enables them to produce more cytokines, and in some cases also more clonal expansion, than naïve cells at lower dose of antigen (1, 2). Further, they produce and release cytokines and cytolytic effectors quickly in response to antigen (3, 4). Memory T cells also have reduced requirement for costimulation (5) and show multi-functionality typically absent in freshly primed T cells (6). All these properties contribute to enhanced protective function of memory T cells along with their increased precursor frequency.

T-cell intrinsic mechanisms responsible for these enhanced functionalities of memory T cells have not been clearly established. There are multiple lines of evidence to implicate epigenetic mechanisms in rapid and increased production of effector molecules (7, 8). Early reports also implicated enhanced TCR signalling (2), however all of the recent studies in fact point to diminished output of TCR signalling in memory cells (9–11). Similarly, oligomeric clusters of TCR seen in antigen-experienced T cells have been implicated in their enhanced production of cytokines (12). But, recent reports argue that TCR is randomly distributed on the plasma membrane of antigen-experienced cells in monomeric state (13) and that they exclusively drive antigen recognition (14). An immediate outcome of antigen recognition and supra-threshold TCR signalling in rapidly migrating, poorly adhesive T cells is the formation of an immature immunological synapse (IS) (15). The immature IS refers to a phase of interaction lasting a few minutes characterized by rapid spreading and adhesion (16, 17). This transient but committed phase further develops into a stable, mature IS lasting over an hour or a motile immunological kinapse (IK) that can nonetheless result in durable interaction (16, 18, 19). We define the capability to form an immature IS as ‘synapse propensity’ and consider it as an intrinsic property of a T cell. Synapse propensity ultimately determines the fraction of precursor cells participating in the response. Therefore, it is appealing to consider the possibility that memory T cells have enhanced synapse propensity compared to the naïve cells. However, diminished output of TCR signalling in memory T cells may also result in lower synapse propensity. We measured synapse propensity of naïve and memory cells from different T cell subsets by multiple approaches. Among the subsets that we have examined, only the human memory CD8 T (CD8^+^ hTm) cells exhibit appreciably higher synapse propensity than naïve counterparts. We have explored the consequence of higher synapse propensity in an ex vivo setting that mimics spatially limiting antigen presentation. We find that higher synapse propensity of CD8^+^ hTm cells gives them a competitive advantage at the expense of naïve T cells.

## Materials and Methods

Most of the experimental and analysis procedures used here have previously been explained in detail (19).

### Isolation of Resting T cell subsets

Non-clinical and de-identified leukapheresis products from donor blood were used as a source of resting human T cells. This was exempt from IRB review at the NYU Medical Center and was approved by National Health Service at the University of Oxford (REC 11/H0711/7). Resting human T cell subsets were isolated using negative selection kits (Stemcell Technologies). The NYU Medical Center Institutional Animal Care and Use Committee approved (Protocol 150609-01) experiments involving mice. Naïve (CD44-ve) and Listeria-specific (CD44+ve) memory CD8 T cells were isolated from spleens of 8-12 week old B6 male mice (from National Cancer Institute, Bethesda, MD) 30-40 days after infected with 5000 colony forming units of *Listeria monocytogenes* The relevant populations were isolated by immunofluorescence flow cytometry sorting (BD FACS Aria).

### Preparation of stimulatory surfaces

Labtek 8-well chambers (Nunc) were used for uniformly coated stimulatory surfaces. CCL21 (3 μg/ml, from Peprotech) was coated first, followed by ICAM1 (2 μg/ml, produced inhouse) and OKT3 (varied concentration, from BioXCell) together. Micro-contact printing technology was used to obtain repeating arrays of circular, stimulatory ‘spots’ with immobilized OKT3 (or 2C11), essentially as described previously (19). These repeating ‘spot’ patterns spanned the entire length of channel of the sticky-Slide VI^04^ (Ibidi), so that bulk assays could also be performed from collected cells. The primary difference in this study is that the coverslips for micro-contact printing were cleaned in 30% 7X Cleaning Solution (MP Biomedicals) on a hot plate (set for 205 °C) for 30-40 minutes, followed by extensive rinsing under flowing deionized water. Cleaned and dried coverslips were then baked at 400 °C in a furnace for 10 hours, cooled to room temperature and used for stamping within a week. The stamped coverslips were affixed to the sticky-Slide and the channels were coated sequentially with CCL21 (13.5 μg/ml) and ICAM1 (3 μg/ml). Supported Lipid Bilayers (SLBs) presenting fluorescent-dye conjugated ICAM1 (200 molecules/μm^2^) and UCHT1 Fab` (varied density, produced in-house) were also assembled in sticky-Slide VI^04^ channels.

### Imaging

Cells were imaged using either a Zeiss LSM 510 (40x Plan Neofluar oil immersion objective with 1.3 NA) or an Olympus FluoView FV1200 (30x Super Apochromat silicone oil immersion objective with 1.05 NA) confocal microscope that was enclosed in an environment chamber (at 37°C) and operating under standard settings. Differential Interference Contrast (DIC) micrographs in time-lapse series were used for detecting and tracking cells. Interference Reflection Microscopy (IRM) images were used for ascertaining spreading or attachment based on the dark patches due to destructive interference. In experiments involving naïve and memory subsets together, the cells were differentially labelled with CellTracker dyes (Life Technologies). The location of stimulatory spots was recorded using Alexa Fluor 647 conjugated to the stamped OKT3 or 2C11 antibody. IS formed on SLBs were imaged in samples fixed with 4% formaldehyde in phosphate buffer saline. High-resolution images were acquired on Olympus CellTIRF-4 line system using 150x 1.45 NA oil immersion objective in Total Internal Reflection Fluorescence (TIRF) mode.

### Image analysis

The time-lapse images were pre-processed in ImageJ as described previously (19). Tracking from DIC, and quantification of associated information on tracked cells from reflection and fluorescence channels, was conducted using TIAM, a MATLAB based toolset that we previously developed (20). Bespoke functions and scripts were written in MATLAB for the calculation of various correlates of synapse propensity from the output of TIAM. These functions and scripts and additional information on image processing are made available on Github (https://github.com/uvmayya/synapsePropensity).

### Quantification of fraction of cells forming IS or IK on SLB

Those fixed cells with >50% of the area under attachment as per the IRM channel and having a ‘circle-like’ attachment footprint were counted as having formed IS or IK. Cells with no or <50% area under attachment or discontinuous or highly elongated or highly irregular or fan-shaped attachment footprint were disregarded while counting cells with IS or IK. These criteria rule out highly motile cells in low adhesive state. These criteria also ensure that both IS and IK are counted together without distinguishing them and thus allow for counting all cells that have passed through the phase of immature IS. Manual counts were corrected for less than 100% purity of naïve and memory subsets, as assessed by flow cytometry, and also for deviation from 1:1 ratio when mixed together after differential labelling. Finally, fraction of cells is obtained based on the expected number of all cells in the field when introduced at the same density (1.5 million/ml). This was done to prevent overestimation of fraction of cells forming IS or IK, as non-attached cells tend to get washed away (or displaced) during fixation and subsequent washes.

### Quantification of fraction of cells forming IS or IK on coated surfaces

Attached segments of tracks from live imaging were selected first, for which we calculated arrest coefficient with the threshold speed of 3 μm/min, implying arrest or deceleration due to IS or IK formation. Any attached segment with arrest coefficient of >0.4 was selected to have led to IS or IK formation in the cell for a significant period during the observation. These relaxed criteria for deceleration allow for inclusion of all cells that have passed through the phase of immature IS. Average number of all cells in the field over the duration of the time-lapse (typically ~50) was used to get the fraction of cells with IS or IK.

### Calculation of on-rate of attachment, encounter rate and arrest efficiency on stimulatory spots

DIC images with binary masks, both for attachment and location of spots, were used to select cells arrested and attached to spots. Plotting number of arrested cells on spots over time gives the attachment curve. The slope of attachment curve over the first ~40 minutes was used for naïve cells whereas data for the first ~15 minutes was used for the rate of attachment (on-rate) of memory cells. Many intervening data points were often removed in the case of naïve cells as there were no additional arrest events and this improved the regression coefficient (R^2^) of linear fitting to be >0.85. The slopes were normalized as per the relative number of cells to spots. The normalization also assumed 1 cell arresting on 10 μm spots and 4 cells arresting on 20 μm spots, which is the typical scenario. Non-responding cells were manually identified and disregarded during normalization, based on non-polarized or shrunken appearance and Brownian motion.

Portions of tracks wherein a cell is on a spot, based on fluorescence value associated with the track-positions, are identified as encounter events. Total number of such ‘clipped’ tracks gives the total number of encounter events. All encounter events during the first 90-120 minutes for naïve cells and 45-60 minutes for memory cells were tallied and divided by the duration of time to calculate encounter rate. Arrest efficiency is defined as the number of separate arrest events on spots over all encounters (transient and durable) with spots for all the cells in the field within a certain period of time. Apart from the presence of a spot beneath a cell for at least 4 minutes, additional criterion of attachment is also included to define arrest events on spots.

### Naïve T cell activation in a competitive setting

Each Ibidi sticky-Slide VI^04^ channel contains ~63,000 spots that are 10 μm wide and 30 μm apart and ~22,800 spots that are 20 μm wide and 50 μm apart. Typically, one cell arrests on spots 10 μm in diameter with ~15% of the spots accommodating two cells. Owing to 4-fold larger area, 4-5 cells arrest on spots 20 μm in diameter. In order to create a competitive setting we used ~120,000 cells on 10 μm spots and ~180,000 cells on 20 μm spots. This is referred to as ‘1x’ number of cells in the Results section. After 10-12 hours, the cells were collected using ice-cold PBS containing 0.5/ BSA and 2mM EDTA. Activation was assessed based on flow cytometry after staining with anti-CD45RO, CD69 and CD62L antibodies. Our isolation procedure for memory cells removes EMRA (<u>E</u>ffector <u>M</u>emory that are CD45<u>RA</u>+ve) cells but retains central and effector memory cells. This total pool of memory cells is used in the competition experiments with naïve cells. In the case of humans, this is physiologically relevant as both spleen and lymph nodes contain central and effector memory cells at comparable frequency to that in the peripheral blood, with considerably lower frequency of EMRA cells in lymph nodes (21).

### Statistical analyses

Statistical tests were performed using Prism (Graphpad). Statistical significance of difference in values, where in a pair of values represent the T cell subsets of a donor, was calculated by paired t-test. P-value from two-tailed tests are denoted as follows in the figures: * for p<0.05, ** for p < 0.01, *** for p < 0.001, and **** for p< 0.0001. If the pairing itself was found to be significant (i.e. p < 0.05), the asterisk rating above the plot are given within parentheses.

## Results

### Human memory CD8 T cells have high synapse propensity

We first assessed synapse propensity CD8^+^ hTm and naive cells on SLBs presenting freely mobile ICAM1 and UCHT1 Fab’. Typically, we use 30 molecules/μm^2^ of UCHT1 Fab’ for imaging synapse formation in individual cells at high resolution in TIRF mode. However, we reasoned that difference in synapse propensity can be appreciated better at very low ligand density (0.3 molecules/μm^2^). Fraction of cells that have passed through the phase of immature IS represents synapse propensity in this context of spatially uniform ligands as this does not typically result in a competitive setting. Since immature IS represents a transient phase committed to durable interaction, we relied on identifying cells that have formed IS or IK after a period of interaction with ligands on SLBs. For unbiased and statistically rigorous counting of cells with IS or IK, we used differentially labelled mixture of naïve and memory cells, fixed them after 30 minutes of interaction with ligands on SLBs and imaged multiple larger fields afforded by lower magnification and lower numerical aperture (Figure 1a). We used attachment areas recorded by Interference Reflection Microscopy (IRM) to decide whether a cell has formed a IS or IK, as visualizing formation of central Supramolecular Activation Cluster (cSMAC) under these settings was not practical. More memory CD8 T cells had attachment footprints typical of cells with IS or IK (Figure 1a, see Methods for scoring criteria). In order to ascertain that these cells were indeed forming bona fide IS or IK, we imaged the same preparations at high resolution in TIRF mode (Figure 1b). cSMAC formation, as assessed by centralized accumulation of UCHT1 Fab’, happened efficiently over 100-fold range of ligand density. When examined over many cells, we found that 16 out of 22 cells that would have been scored as having formed IS or IK based on attachment footprint had a typical cSMAC even at 0.3 molecules/μm^2^ of UCHT1 Fab’ (Supplementary Figure 1). All 22 cells had substantially enriched UCHT1 Fab’ and ICAM1 in the interface compared to the surrounding (Figure 1c). This confirmed that all CD8^+^ hTm cells with attachment footprint typical of IS had indeed formed IS. We tallied the fraction of cells forming IS or IK on SLBs and expectedly found that synapse propensity increases with increasing surface density of UCHT1 Fab’ for all subsets examined (Figure 1d and 1e). Overall, the memory cells have higher synapse propensity than naïve counterparts. The difference in synapse propensity between naïve and memory cells is more striking in the CD8 subset and CD8^+^ hTm cells have appreciably higher synapse propensity than the CD4^+^ hTm cells on SLBs presenting freely mobile ligands.

**Figure 1:**
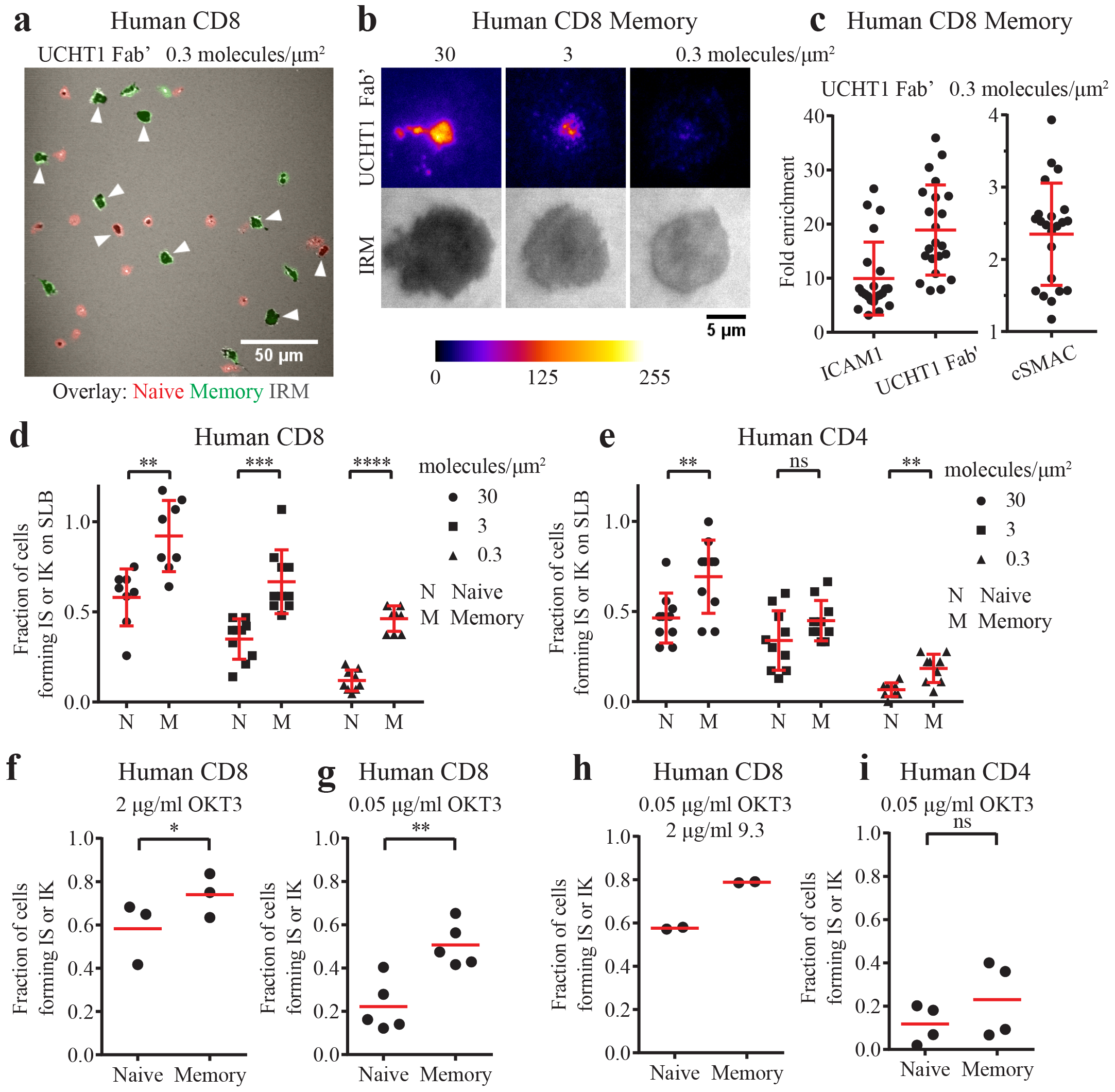
CD8^+^ hTm cells have high synapse propensity on uniformly presented stimulatory surfaces. a) CD8^+^ hTm (in green) and naïve (in red) cells forming IS or IK on SLBs presenting 0.3 molecules/μm^2^ of UCHT1 Fab’. The dark patches in the fluorescence micrograph, in fixed cells captured by confocal microscopy (30x, 1.05 NA), are the result of overlay of the IRM image signifying attachment of the cell. Attachment along with additional criteria (see Methods) are used to count cells with IS or IK. Such cells are highlighted with a white triangle in this example. More CD8^+^ hTm cells pass through the phase of immature IS compared to naïve cells and then form IS or IK. b) cSMAC formation by CD8^+^ hTm cells at 30, 3 and 0.3 molecules/μm^2^ of UCHT1 Fab’ as visualized in fixed cells by TIRF microscopy (150x, 1.45 NA). Centralized accumulation of UCHT1 Fab’ is a result of cSMAC formation. The cells are able to form cSMAC even at very low density of UCHT1 Fab’. c) Enrichment of ligands due to continued accumulation in the interface of those cells with IRM footprint that is typical of IS or IK formation. Fold enrichment is the mean signal intensity in the interface relative to mean signal intensity in the surrounding region outside of the cell (i.e. background). For enrichment within cSMAC, mean intensity in the most intense region is expressed relative to that of the entire interface. Each data-point represents a cell. Mean and standard deviation are shown in red. d and e) Fraction of naïve and memory cells, in the human CD8 (d) and CD4 (e) subsets, forming IS or IK on SLBs at varying density of UCHT1 Fab’. Due to the use of expected number of cells in the field as the denominator, some data-points have a value of >1. ~30 cells per field would represent 100/ of the cells forming synapse under our experimental condition. 8-10 fields per condition and subset were imaged and plotted. The data shown is representative of two independent experiments. f to i) Fraction of human naïve and memory T cells forming IS or IK on uniformly coated surface with immobilized CCL21, ICAM1 and OKT3. Note the differences in OKT3 concentration (f vs. g) used for coating, presence of anti-CD28 antibody 9.3 (in h) and the data for the CD8 vs. CD4 subset (in i). The cells were motile because of the presence of immobilized CCL21. Time-lapse data (1.5-2 hours) was used to determine the number of cells forming IS or IK based on scoring for attachment and deceleration (see Methods). Each data point represents a separate donor and independent experiment. Mean value is shown in red. See Methods section for interpreting statistical significance.

We next assessed synapse propensity of CD8^+^ hTm and naïve cells on uniformly coated surfaces presenting immobilized CCL21, ICAM1 and OKT3. Effector memory cells are thought of as being negative for expression of CCR7, the receptor for the chemokine CCL21 (22). However, reagents to detect human CCR7 have likely improved in the two decades since. Therefore, we first compared the expression of CCR7 and responsiveness to CCL21 in naïve and memory cells (Supplementary Figure 2). We noted that both central and effector memory CD8 T cells express sufficient CCR7 to show robust chemokinesis on immobilized CCL21. In fact, the speed of CCL21-induced motility is higher for central and effector memory cells than naïve cells. Thus, we can rule out the possibility of lack of motility indirectly promoting synapse propensity in memory cells. We proceeded to score the fraction of cells forming IS or IK on uniformly coated surfaces by live imaging based both on attachment and deceleration (or arrest). As in the case of SLBs, synapse propensity reduced with less OKT3, but the difference between naïve and memory CD8 T cells was more pronounced with less OKT3 (Figure 1f and 1g and Supplementary Video 1). High synapse propensity of CD8^+^ hTm cells was maintained even with costimulation from immobilized anti-CD28 (9.3) antibody (Figure 1h). No significant difference in synapse propensity was observed between naïve and memory human CD4 cells (Figure 1i). Overall, we conclude that CD8^+^ hTm cells have high synapse propensity on uniformly presented stimulatory surfaces.

In vivo, both under the settings of priming in secondary lymphoid organs and during the early phase of recall response in peripheral tissues, antigen can be sparse and presented on few dispersed antigen-presenting cells in a spatially limiting manner (23). We have recapitulated this scenario ex vivo using the micro-contact printing technology by creating stimulatory spots of OKT3 with pervasive ICAM1 and CCL21 (Figure 2a) (19). CD8^+^ hTm cells rapidly attached and arrested on the stimulatory spots when compared to the naïve CD8 T cells (Figure 2b and Supplementary Video 2). The rate of arrest is the best measure of synapse propensity on spatially limiting stimulatory spots, as this represents a competitive setting. For an accurate measure of on-rate of arrest on the spots, we took the slope of the initial part of the attachment curve. We found that memory cells were capable of arresting on 10 μm spots at 7.1-fold and on 20 μm spots at 5.4-fold faster rate than naïve cells (Figure 2c and 2d). Based on the principles of bimolecular reaction kinetics, on-rate of arrest should be the product of rate of encounter with spots and arrest efficiency. Encounter rate quantifies the rate at which cells search and locate the stimulatory spots and arrest efficiency quantifies the ability to attach and arrest upon locating a spot. Encounter rate of memory cells is ~1.5 fold higher than that of naïve cells Figure 2e and 2f). This is in agreement with ~1.6 fold increased speed of memory cells on immobilized CCL21 (Supplementary Figure 2c). Arrest efficiency of CD8^+^ hTm cells is 5.1 fold higher on 10 μm spots and 3.5 fold higher on 20 μm spots than that of naïve CD8 T cells (Figure 2g and 2h). Roughly one in ten encounters leads to arrest on the spots in the case of naïve CD8 T cells whereas ~40% of the encounters lead to arrest in the case of CD8^+^ hTm cells. As expected, multiplying these two parameters gives back the same on-rate of attachment as the value initially measured by mathematically independent approach. Consistent with the observation on uniform stimulatory surface, there is no difference in synapse propensity of human CD4 naïve and memory cells on stimulatory spots, with both having low values (Supplementary Figure 3a-3c). Surprisingly, even the mouse CD8 naïve and memory cells do not show any significant difference in synapse propensity on stimulatory spots (Supplementary Figure 3d-3e). Thus, CD8^+^ hTm cells uniquely show very high synapse propensity both on uniform stimulatory surface and spatially limiting stimulatory spots.

**Figure 2:**
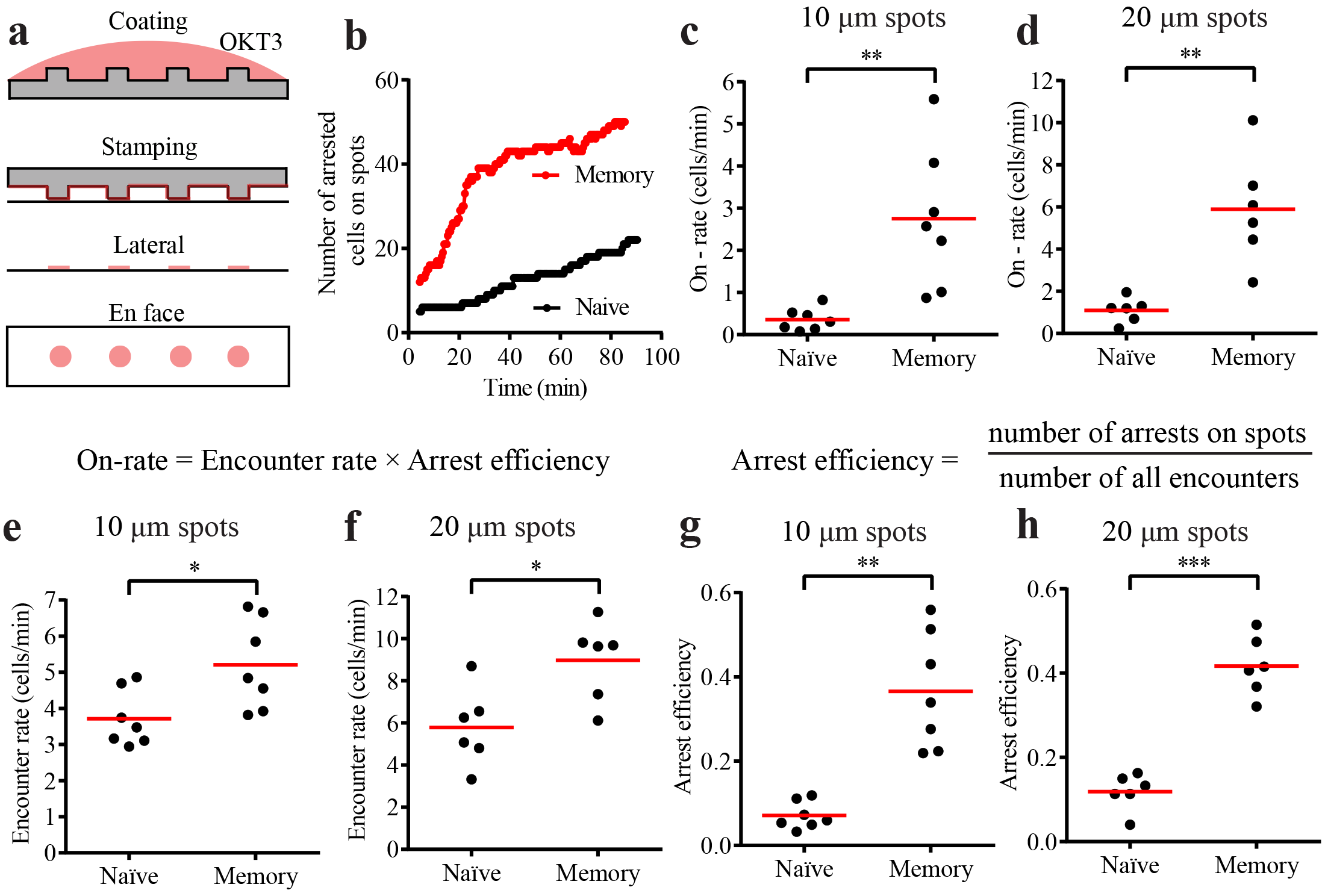
CD8^+^ hTm cells have high synapse propensity on spatially limiting stimulatory spots. a) Schematic of the micro-contact printing procedure. Antibody is first adsorbed to polydimethylsiloxane (PDMS) casts that have short pillars with flat tops that will generate the spots. When the cast is laid on the coverglass, some of the antibody gets transferred from the PDMS surface onto the coverglass. However, this only happens at the top of the pillars. CCL21 and ICAM1 are then adsorbed uniformly across the surface from solution. b) Number of CD8^+^ hTm and naive cells arresting on 10 μm spots that are 30 μm apart over 90 minutes. 64 spots and typically at least the same number of cells were present in the field in such experiments. c and d) Rate of attachment and arrest (on-rate) of human CD8 T cells on 10 μm spots that are 30 μm apart (in c) and 20 μm spots that are 50 μm apart (in d). The on-rate represents the values for the imaging field that is 225 μm by 225 μm in size, hence the absence of mm^-2^ in the unit. e and f) Rate of encounter of human CD8 T cells with 10 μm (in e) and 20 μm (in f) spots. Again, the values given are for the specific imaging field. g and h) Arrest efficiency of human CD8 T cells on 10 μm (in g) or 20 μm (in h) spots. Both encounter rate and arrest efficiency contribute to very high on-rate of attachment of CD8^+^ hTm compared to the naïve cells. Each data point represents a separate donor and independent experiment in each plot. Mean value is shown in red. See Methods section for interpreting statistical significance.

### High synapse propensity of human memory CD8 T cells confers competitive advantage in recruitment to the stimulatory spots and can suppress naïve cell activation

We reasoned that in a competitive setting of both cell types present in equal numbers, CD8^+^ hTm cells will prevent naïve cells from arresting on the stimulatory spots. Expectedly, memory cells rapidly arrested on spots over the first 2 hours leaving the naïve cells to explore area outside the spots (Supplementary Video 3). The enrichment of CD8^+^ hTm cells, as per the relative number of cells on spots after 2 hours, approximately matched their fold-increase in synapse propensity, i.e. on-rate of attachment, both on 10 and 20 μm spots (Figure 3a and 3b). Again, as predicted by lack of difference in on-rate of attachment (Supplementary Figure 3a), there was equal sharing of spots between naïve and memory human CD4 T cells (Figure 3c). There was a marginal increase in on-rate of murine naïve CD8 T cells (Supplementary Figure 3d), which explains the slight enrichment of naïve cells over memory cells on spots deposited with 2C11 (Figure 3d).

**Figure 3:**
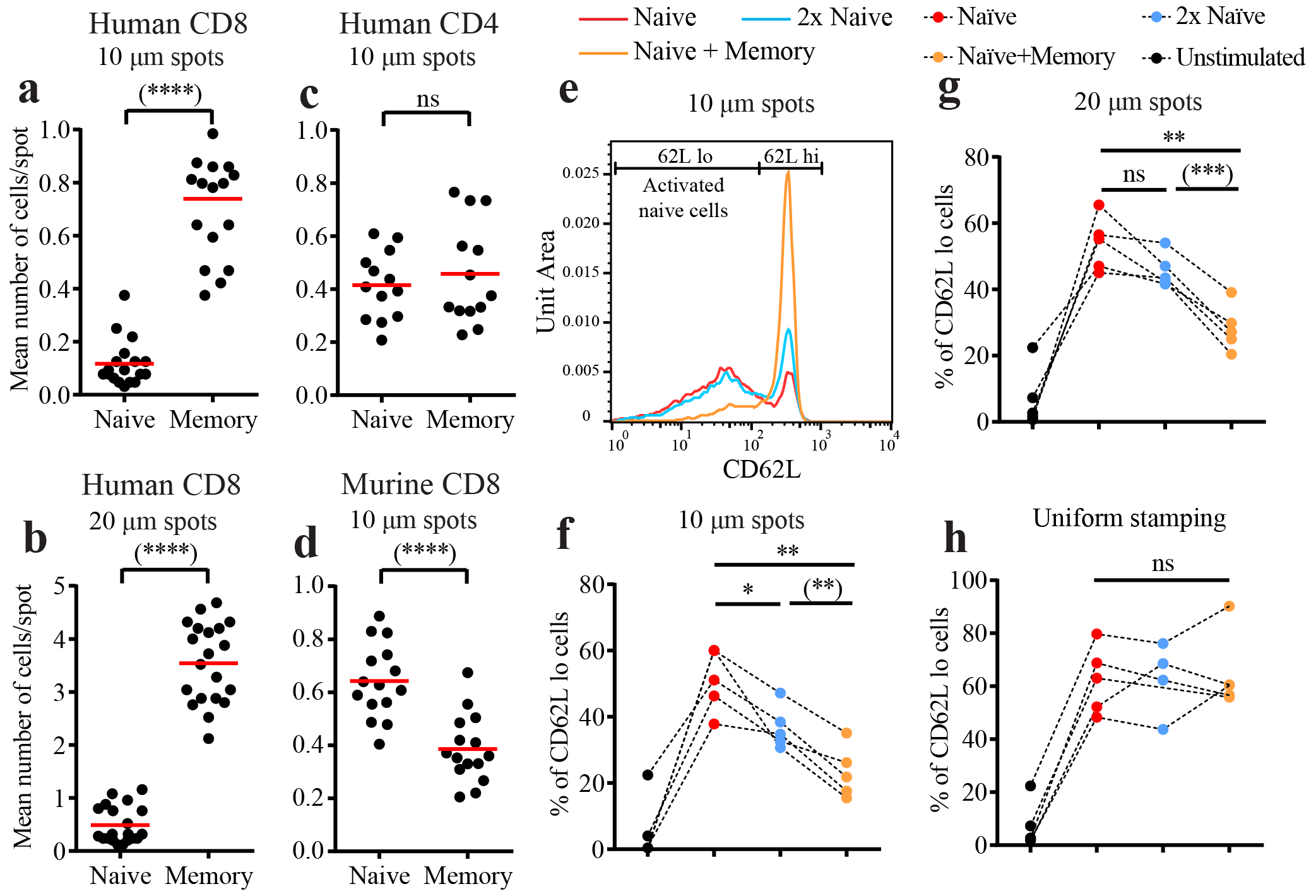
CD8^+^ hTm cells compete out naïve T cells by preventing access to stimulatory spots due to their high synapse propensity. a to d) 1x number (see Methods for details) of differentially labelled naïve and memory cells were pooled and introduced into the channel with stimulatory spots. After 2 hours, multiple fields along the channel were imaged for multiple donors. Mean number of cells arrested per spot in the field is plotted for each field imaged. Type of spot configuration and T cell subset examined is denoted at the top and in category names of the dot plots (Human CD8 T cells in a and b, Human CD4 T cells in c, and Murine CD8 T cells in d). Mean value is shown in red. e) Activation of human naïve CD8 T cells assessed after 10-12 hours of interaction with 10 μm wide stimulatory spots in a competitive setting by staining for CD62L. Activated cells have lower levels of CD62L due to gradual proteolytic shedding during continuous TCR signalling. Note that the histograms have the same number of naïve cells represented. Having 2x number of naïve cells is a ‘control’ case (blue) to compare against the scenario wherein memory cells are also present (orange). Note that the presence of memory cells reduces not only the % of naïve cells that get activated but also the extent of shedding of CD62L. This implies that memory cells reduce TCR signalling even among the activated naïve T cells. f to h) % of activated naïve CD8 T cells, as measured by the % of cells with lower CD62L, under different competitive settings on 10 μm spots (in f), 20 μm spots (in g), and on uniformly stamped surface (in h). Lines that join the dots for each competitive setting represent the same donor. Data points for each competitive setting are color-coded (see top of panel g). 1x number of cells on 20 μm spots is considerably higher (see Methods). Channels with uniformly stamped surface received the same number of cells as the channels with 10 μm spots under the corresponding competitive settings. See Methods section for interpreting statistical significance.

We next investigated how synapse propensity of CD8^+^ hTm cells impacts activation of naïve T cells in a competitive setting. We had earlier shown that stimulatory spots with anti-CD3 can robustly prime human naïve CD8 T cells to undergo full activation and multiple rounds of cell division (19). Here, we looked at activation after 10-12 hours under a competitive setting to begin with, as described in the Methods section. Substantial fraction of naïve cells proteolytically shed CD62L due to continuous TCR signalling (Figure 3e). We have shown before that naïve cells with low CD62L after 12 hours are activated cells, as indicated by robust CD69 upregulation. With twice the number of naïve cells, the fraction of activated cells expectedly reduces (Figure 3e). However, with the presence of equal number of memory CD8 T cells, the fraction of naïve cells activated reduces even further (Figure 3e). The same effect of suppression of naïve cell activation by CD8^+^ hTm cells is observed across multiple donors and both on 10 μm and 20 μm spots (Figure 3f and 3g). However, we do not see any suppression of naïve cell activation in the presence of memory cells on uniformly stamped surface wherein immobilized anti-CD3 is available across the entire surface (Figure 3h). This implies that memory cells do not secrete any suppressive soluble factor nor do they participate in any contact-dependent mechanisms of suppression. The suppression of naïve cell activation by memory cells is essentially because of higher synapse propensity of memory cells preventing access to the stimulatory spots. However, the extent of suppression seen after 10-12 hours is considerably less than what one would have anticipated based on the competitive advantage seen after 2 hours (Figure 3a vs. Figure 3f, for example). This is because of reduced durability (half-life) of interaction of CD8^+^ hTm cells on the stimulatory spots (19), allowing naïve cells to gradually get access to the spots over longer periods of time. In fact, the extent of suppression, when compared against twice the number of naïve cells, is in agreement with steady state behaviour predicted by the combined effect of on-rate (i.e. synapse propensity) and off-rate (i.e. inverse of durability). Ultimately, the memory cells have ~2 fold competitive advantage over naïve CD8 T cells, resulting from ~7 fold higher synapse propensity (Figure 2) and ~3.5 fold lower durability (19) on the 10 μm stimulatory spots. Overall, high synapse propensity of CD8^+^ hTm cells allows them to compete out naïve cells from getting activated.

### Higher surface levels of LFA1 contributes to increased synapse propensity of CD8^+^ hTm cells

Both CD8^+^ hTm and naïve cells have the same surface levels of CD3 (Supplementary Figure 4a and 4b), ruling out that being a potential player in increased synapse propensity. However, memory cells express ~2.5 fold more LFA1 (Figure 4a, Supplementary Figure 4c) (24). We sought to find out if increased expression of LFA1 contributed to increased synapse propensity of CD8^+^ hTm cells. Based on fitting the binding curve for the blocking antibody TS1/18 with Scatchard equation, one can determine the concentration of antibody needed to effectively bring down the level of functional LFA1 in memory cells to what is found on naïve cells (Figure 4b). When the level of LFA1 was brought down to that seen in naïve cells from the same donor using the blocking antibody TS1/18, memory cells exhibited lower synapse propensity both on uniformly coated surface (Figure 4c and Supplementary Video 4) and on stimulatory spots (Figure 4d). Arrest efficiency rather than on-rate of arrest is reported for stimulatory spots as there was no change in the rate of encounter with the spots due to reduced levels of functional LFA1 in memory cells. We confirmed the finding by independent means by using a small molecular antagonist of LFA1, BIRT377. Even with modest concentration of 0.3 μM BIRT377 (IC50 in adhesion of SKW3 cells to immobilized ICAM1 was 2.6 μM), synapse propensity of memory CD8 T cells was reduced both on uniformly coated surface (Figure 4e) and on stimulatory spots (Figure 4f and Supplementary Video 5). In conclusion, higher surface levels of LFA1 on CD8^+^ hTm cells contributes to their high synapse propensity.

**Figure 4:**
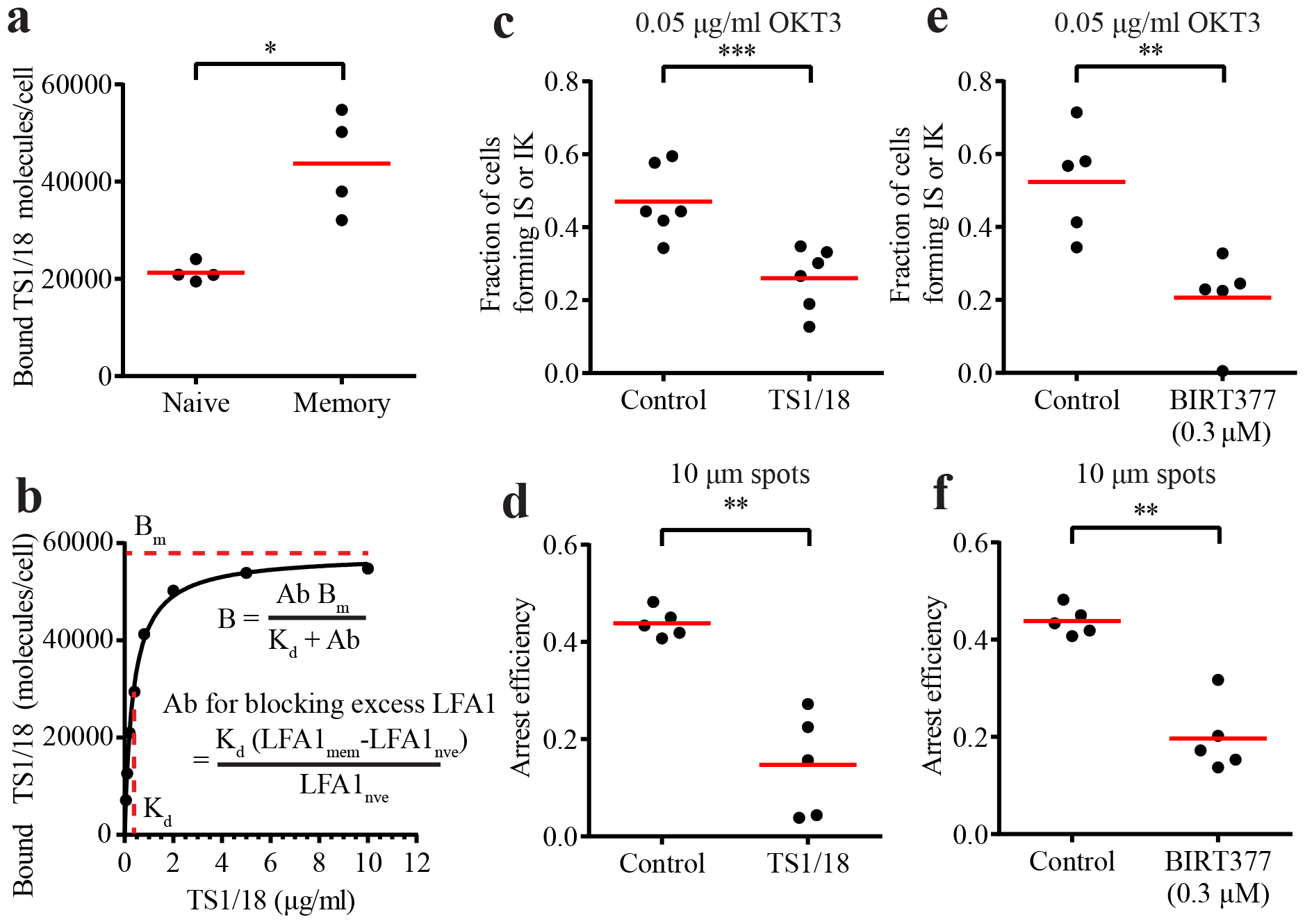
Higher surface expression of LFA1 in CD8^+^ hTm cells contributes to high synapse propensity. a) CD8^+^ hTm cells have more surface LFA1 than naïve cells. Surface expression of LFA1 was estimated by labelling the cells with TS1/18, a blocking antibody against the β subunit of LFA1, under saturating conditions (10 μg/ml) such that virtually all LFA1 is bound. Absolute numbers of antibody molecules comes from the use of standardized beads with known number of fluorochrome equivalents in flow cytometry measurements. b) Binding of TS1/18 to memory cells as a function of concentration. Based on the estimate of Kd (dissociation constant) of binding from curve fitting (Scatchard equation, on top), it is possible to determine the concentration of TS1/18 needed to block the excess number of LFA1 molecules and thereby bring down the effective number of functional LFA1 molecules to what is seen in naïve cells. The plot is a representative example for illustration. c and d) Synapse propensity of CD8^+^ hTm cells is reduced after blocking excess LFA1 molecules with TS1/18, both on uniformly coated surface (c) and on 10 μm-wide stimulatory spots (d). e and f) Synapse propensity of CD8^+^ hTm cells is reduced after inhibiting LFA1 with BIRT377 both on uniformly coated surface (e) and on 10 μm-wide stimulatory spots (f). The synapse propensity measurements were carried out as described in earlier figures. Cells were treated with TS1/18 or BIRT377 for 10-20 minutes before initiating imaging. Each data point represents a separate donor and independent experiment in each plot, except in b. Mean value is shown in red. See Methods section for interpreting statistical significance.

## Discussion

CD8^+^ hTm cells, uniquely, have a ~7-fold higher synapse propensity compared to the naïve counterparts (Figure 2). This means they have 7-fold higher probability to switch from low-adhesion scanning motility to high-adhesion immature IS when they come in contact with antigen. In a recent study, we also showed that CD8^+^ hTm predominantly formed stable mature synapses, whereas naïve CD8^+^ T cells predominantly formed motile kinpses that were ~3.5 fold more durable on the stimulatory spots (19). Synapse propensity, IS vs. IK ratio and durability are all distinct cell-intrinsic parameters of interaction with antigen. Synapse propensity and durability determine competition in a linear manner such that CD8^+^ hTm have a ~2 fold-advantage over naïve cells while competing for antigen (Figure 3).

How and why do the CD8^+^ hTm cells have high synapse propensity? With regard to ‘how’, we have shown that increased expression of LFA1 contributes to high synapse propensity of CD8^+^ hTm cells (Figure 4). LFA1 cannot be sufficient for high synapse propensity as all murine and human CD4 and CD8 memory cell subsets express more than their naïve counterparts. We think of LFA1 as more of a handle to access the intrinsic responsiveness of human memory CD8 T cells in our reductionist ex vivo system. This intrinsic responsiveness ensures appreciably more number of productive TCR triggering events within a minute thus increasing the probability of calcium influx and synapse formation (15, 25). What then determines the intrinsic responsiveness that leads to high synapse propensity? Differential expression analysis of publicly available microarray datasets did not reveal anything unique about CD8^+^ hTm cells when compared against both CD4+ hTm and murine memory CD8 T cells. This suggests that responsiveness is an emergent property of CD8^+^ hTm cells.

Why do the CD8^+^ hTm cells have high synapse propensity? High synapse propensity should naturally increase the fraction of precursors recruited to the response. It should also increase the synchrony in the response as majority of the cells rapidly form IS. Both of these attributes enhance the protective functionalities of memory cells. However, it is not clear why the property is unique to CD8^+^ hTm cells and absent in other memory subsets examined. The competitive advantage of CD8^+^ hTm cells at the expense of naïve CD8 T cells (Figure 3) provides an important clue to this conundrum. It is known that maintenance of naïve cell pool is important through the life time of the host, and especially critical to the health of aged individuals (26). Reduction in naïve cell number and repertoire, and especially that of naïve CD8 T cells, is a hallmark of human immune aging (27). In mice, thymic output throughout the lifespan ensures constant supply of naïve cells (28). In humans, thymic output is negligible to absent in adult life and homeostatic proliferation maintains the naïve cell pool (28). The CD8^+^ hTm cells, by competing out the naïve cells during the current response, can help preserve the repertoire and number of naïve cells for when they are absolutely needed. Increased cross-reactivity, likely afforded by considerably longer peptides binding to class II MHC, ensures abundance of pre-existing cross-reactive human memory CD4 cells (29, 30). Therefore, additional means are not necessary to help preserve the number of naïve CD4 cells in humans. This also means that high synapse propensity in CD8^+^ hTm cells can promote cross-reactivity. Consistent with this corollary, original antigenic sin, a phenomenon wherein recruitment of naïve cells against new variant epitopes is suppressed by memory cells against the original epitopes, occurs only in human CD8 T cells (31, 32) but not in mice (33). Whether this phenomenon is solely driven by the higher synapse propensity or by other factors remains to be answered.

## Acknowledgement

We thank staff members of Light Microscopy and Flow Cytometry Core Facilities at the NYU Medical Center for their technical support, A. Gondarenko for help with e-beam lithography, S. Valvo, J. Afrose and H. Rada for key bilayer reagents and members of Dustin and Kam labs for discussions. A special thanks also to D. Fooksman for initially suggesting competition among cells as a potential scenario where synapse propensity could have an impact. Finally, we thank Boehringer Ingelheim Pharmaceuticals for providing access to BIRT377.

## Abbreviations used

IS: Immunological Synapse
IK: Immunological Kinapse
CD8^+^ hTm cells: human memory CD8 T cells
SLB: Supported Lipid Bilayer
TIRF: Total Internal Reflection Fluorescence
IRM: Interference Reflection Microscopy
cSMAC: central Supramolecular Activation Cluster

