## supplementary figures and video legends for "Synapse propensity of human memory CD8 T cells confers competitive advantage over naïve counterparts"

### Content

Supplementary Figures 1-4

Legends for supplementary videos 1-5

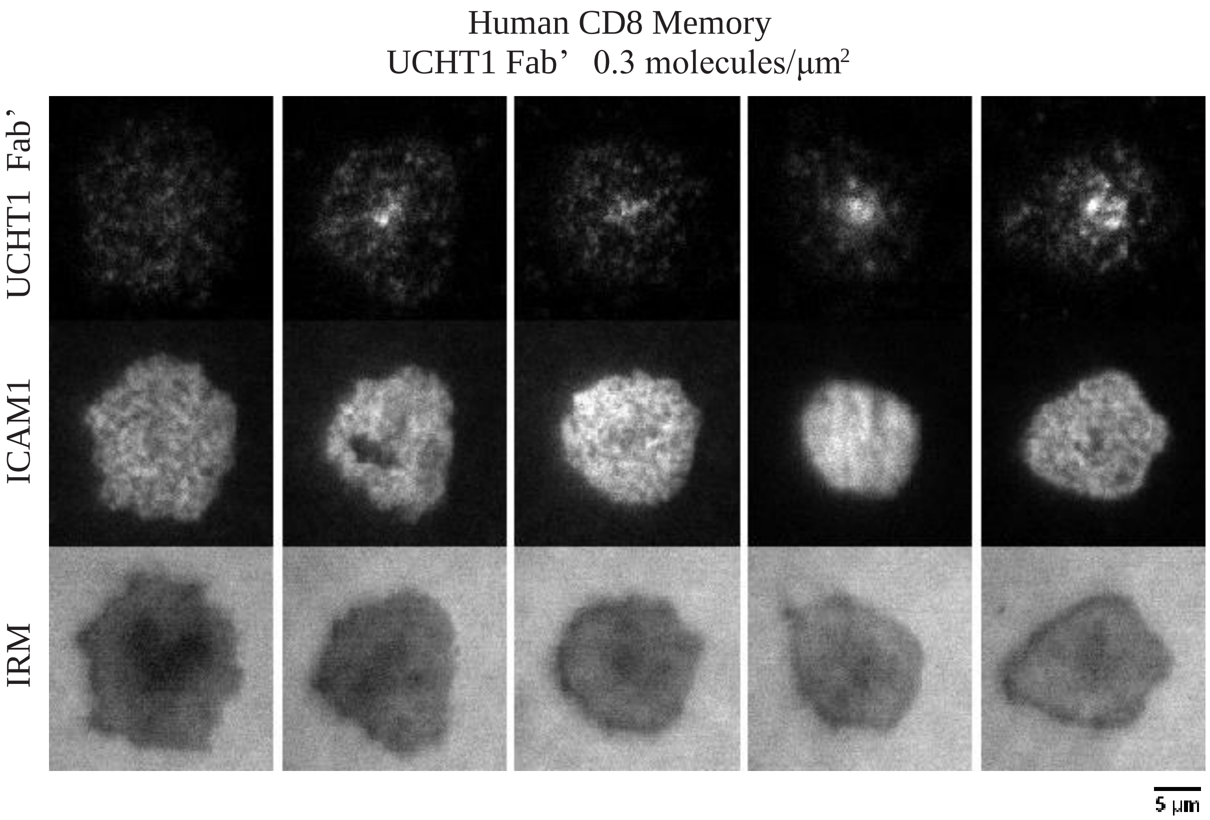


### Supplementary Figure 1

**Supplementary Figure 1:** Synapse formation of $CD8^+$ hTm cells (human memory CD8 T cells) on very low density of UCHT1 Fab' (0.3 molecules/$\mu m^2$) on SLBs. All cells with IRM footprint (bottom row) typical of synapses do show enrichment of ICAM1 (middle row) and UCHT1 Fab' (top row) at the synaptic interface. Most cells also show appreciable central accumulation of UCHT1 Fab' marking cSMAC. Except for the first cell (left most) in these examples, all cells show prominent cSMAC. However, the canonical pSMAC ring is absent as ICAM1 is not excluded at lower densities of UCHT1 Fab'.

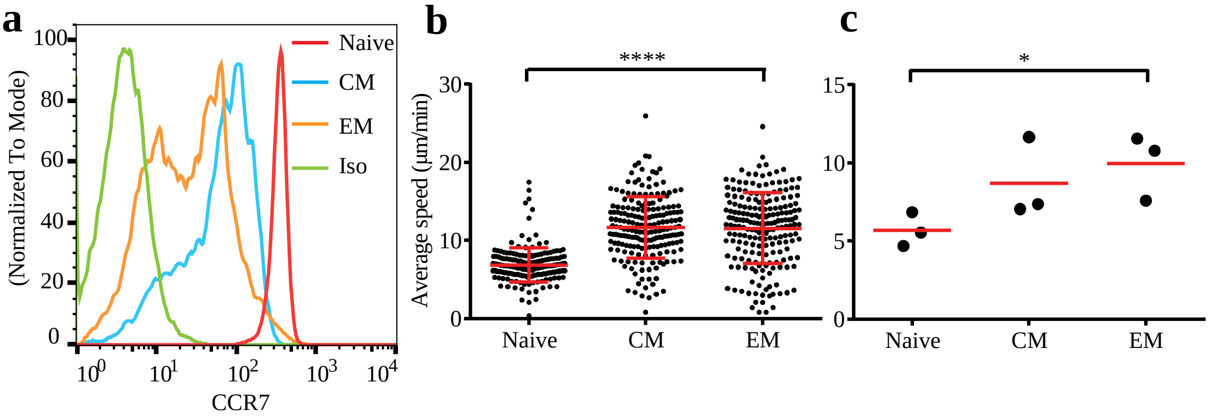

**Supplementary Figure 2:** a) Expression of CCR7 in naïve, central memory (CM) and effector memory (EM) human CD8 T cell subsets probed with CCR7-specific antibody (clone G043H7; Cat. No. 353205 from Biolegend). Representative of two independent experiments/donors. b and c) Average speed of naïve, central memory (CM), and effector memory (EM) human CD8 T cells in response to immobilized CCL21 and ICAM1. When only ICAM1 is present <5% of the EM cells show motility, however along with CCL21 >80% of the cells show motility. Data points in b represent individual cell-tracks, whereas data points in c represent population means from separate donors. Note that central and effector memory cells move faster on immobilized CCL21 despite having reduced surface expression of CCR7.

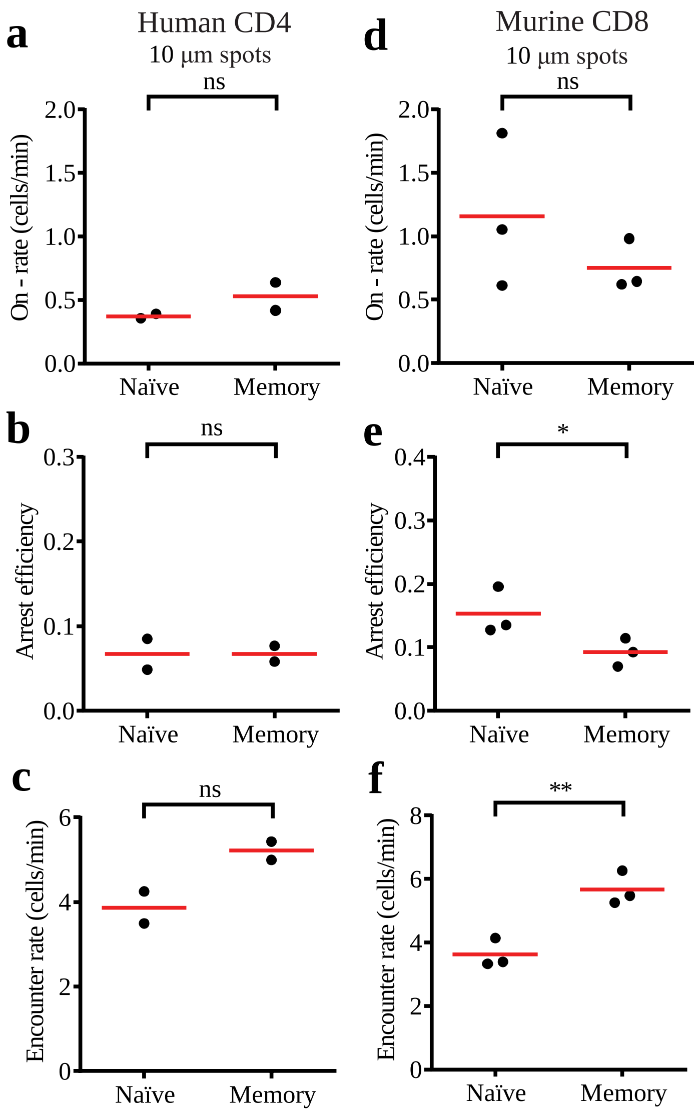


**Supplementary Figure 3:** Synapse propensity of human CD4 (a-c) and murine CD8 (d-f) T cells measured on 10 µm-wide stimulatory spots. Refer to legend for Figure 2 in the main text for details on the calculations for on-rate of arrest, arrest efficiency and encounter rate. There is no difference in any of the measured parameters in the case of human naïve and memory CD4 T cells. While arrest efficiency is higher for murine naïve CD8 T cells, it is offset by higher encounter rate of memory CD8 T cells, ultimately resulting in no significant increase on-rate of attachment and arrest. Each data point represents a separate donor and independent experiment in each plot. Mean value is shown in red. See Methods section for interpreting statistical significance.

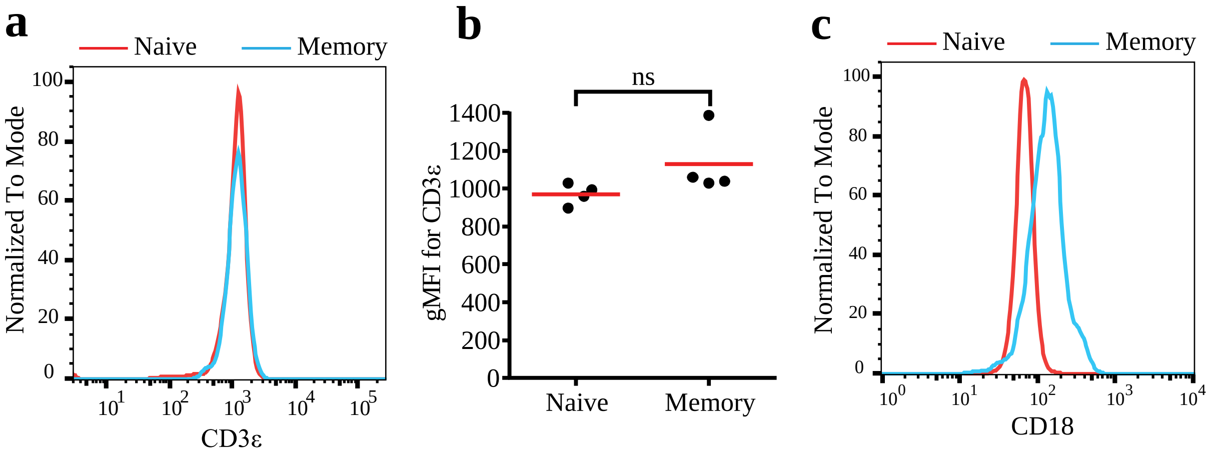


**Supplementary Figure 4:** a and b) Surface expression of CD3ε in human naïve and memory CD8 T cells. Representative histogram from cytometry is shown in a. Geometric mean fluorescence intensity of bound antibody against CD3ε across multiple donors. c) Representative histogram showing increased expression of LFA1 (CD18 is the β subunit in LFA1) in $CD8^+$ hTm cells.

### Legends for Supplementary Videos

**Supplementary Video 1:** Higher synapse propensity of $CD8^+$ hTm cells (green) compared to the naïve cells (red) on uniformly coated surface with CCL21, ICAM1 and low density of OKT3 (0.05 µg/ml used for coating). Equal numbers of differentially labelled naïve and memory cells were introduced into the Nunc well for imaging. The video also contains overlay of the IRM channel for visualize attachment footprints (dark gray). Time-ticker and scale bar are shown. Majority of memory cells attach and arrest (or decelerate) due to synapse formation. Majority of naïve cells only show chemokinesis in response to CCL21. Because of advection in the Nunc wells, most of these motile naïve cells drift from top end of the field to the bottom end.

**Supplementary Video 2:** Higher synapse propensity of $CD8^+$ hTm cells on 10 µm-wide stimulatory spots. Naïve cells are shown on the left and memory cells on the right. The video is an overlay of the spots (magenta) and cells imaged by DIC and IRM. The dark patches show attachment footprints, which happens when cells arrest on the spots. Time-ticker and scale bar are shown. Memory cells arrest on the spots at a faster rate, i.e. the number of spots with arrested memory cells increases rapidly. The field also contains appreciable number of non-responding and dead cells with Brownian motion among the memory cells.

**Supplementary Video 3:** Higher synapse propensity confers competitive advantage to $CD8^+$ hTm cells. Memory (green) cells rapidly arrest and attach on the 10 µm (left side) and 20 µm (right side) spots (in gray). This leaves very little area on the spots for naïve (red) cells to sample and arrest on to, as they have much lower synapse propensity.

**Supplementary Video 4:** Reducing the level of functional LFA1 on $CD8^+$ hTm cells down to that of naïve cells reduces synapse propensity. Here, the antibody TS1/18 against the β subunit is used to block LFA1. Cells are interacting with uniformly coated and immobilized CCL21, ICAM1 and low density of OKT3 (0.05 µg/ml used for coating). The video is an overlay of and cells imaged by DIC and IRM. Most memory cells treated with an isotype control antibody arrest and attach (footprints in dark gray). However far fewer memory cells treated with TS1/18 arrest and attach. The remaining cells are motile due to responsiveness to CCL21, however because of advection in the Nunc wells, they also show drift.

**Supplementary Video 5:** Inhibiting LFA1 using 0.3 µM BIRT377, a small molecule antagonist, reduces the synapse propensity of $CD8^+$ hTm cells on 20 µm stimulatory spots (in magenta). Under the control setting (left side) the memory cells rapidly arrest and attach (footprint shown in dark gray) on the spots. Upon inhibition of LFA1 (right side) very few memory cells arrest and attach on the spots over time.
